# Predicting FDA approvability of small-molecule drugs

**DOI:** 10.1101/2022.10.15.512330

**Authors:** Chih-Han Huang, Justine Hsu, Li-yen Yang, Tsai-Min Chen, Edward S.C. Shih, Ming-Jing Hwang

**Affiliations:** Institute of Biomedical Sciences, Academia Sinica, Taipei 11529, Taiwan; Genome and Systems Biology Degree Program, Academia Sinica and National Taiwan University, Taipei 106319, Taiwan; Graduate Program of Data Science, National Taiwan University and Academia Sinica, Taipei 11529, Taiwan

**Keywords:** Machine learning, Drug discovery, Compound attrition, Physicochemical properties, Molecular fingerprints

## Abstract

A high rate of compound attrition makes drug discovery via conventional methods time-consuming and expensive. Here, we showed that machine learning models can be trained to classify compounds into distinctive groups according to their status in the drug development process, which can significantly reduce the compound attrition rate. Using molecular structure fingerprints and physicochemical properties as input, our models accurately predicted which drug compounds would proceed to trial, with an area under the receiver operating curve (AUC) of 0.94 ± 0.01 (mean ± standard deviation). Our models also identified which drugs in clinical trials would be approved by the US Food and Drug Administration (FDA) to go on the market, with an AUC of 0.73 ± 0.02. The predictive power of our models could reduce the attrition rate of preclinical compounds to enter clinical trials from 65%, as with conventional methods, to 12% (with 92% sensitivity) and the clinical trial failure rate from 80–90% to 29% (with 83% sensitivity). The results largely held in additional tests on new clinical trial compounds and new FDA-approved drugs, as well as on drugs uniquely approved for use in Europe and Japan.

**SIGNIFICANCE STATEMENT:** The odds of developing a drug approved by the US Food and Drug Administration (FDA) are slim, meaning that the vast majority of drug candidates would fail tests for safety and efficacy in the drug discovery process, rendering it highly inefficient and costly. Here, we have developed machine learning models to predict drug compounds worthy of clinical trials with high accuracy, and clinical-trial compounds to receive FDA approval with a much higher success rate than that achieved by the traditional approach. Our computational prediction requires input of only the drug compound’s chemical structure and physicochemical properties. It can help mitigate the long-standing problem of drug discovery.

## INTRODUCTION

It typically takes over 10 years and billions of US dollars for a drug to meet the rigorous standards set by the US Food and Drug Administration (FDA) to approve it for sale.^1^ Most drugs do not meet these standards, with an overall 96% attrition rate which includes a 80-90% failure rate for drugs in clinical trials.^2-4^ Multiple factors contribute to these long-standing issues in the drug discovery process. For example, more than 50% of the compounds that advance to phase III of clinical trials fail due to inefficacy (e.g., weak pharmacokinetic or pharmacodynamic properties), and another 17% fail because of adverse effects.^5^ Even after FDA approval, serious adverse effects, such as heart toxicity, fainting, and death, can occur.^6^ These drugs are usually withdrawn from the market, causing a huge financial loss for pharmaceutical companies.

To alleviate the attrition problem in drug development, the concept of “drug-likeness” was developed to serve as guidelines to increase the likelihood of a compound entering clinical trials and being approved for medicinal use.^7, 8^ Lipinski’s Rule of Five was first proposed to assess a compound’s drug-likeness by a set of four physicochemical properties (PCPs) associated with oral activity derived from clinical drugs.^9^ Other simple rules were subsequently reported, including the Ghose filter,^10^ the Veber filter,^11^ the Rule of Three,^12^ and the Rapid Elimination of Swill (REOS) filter.^13^ These rules are usually based on a compound’s PCPs, such as threshold values for molecular weight, number of hydrogen bond donors and acceptors, number of rotatable bonds, Log P measure of lipophilicity, and so on. PCPs also can help predict absorption, distribution, metabolism, and excretion profiles of small molecules to find desirable compounds for drug discovery.^14^

In addition to PCPs, a drug’s pharmacokinetics and reactivity can be altered by changing its chemical structure. For example, synthetic cathinones are a category of designer drugs with effects similar to amphetamines but with different pharmacokinetic characteristics due to structural modifications (e.g., increased length of the alkyl chain on the α position of methylone resulting in increased plasma concentrations and longer half-life).^15^ Fluoroquinolones, a class of highly effective antibiotics, exhibit bactericidal activities that can be modulated in additive fashion based on the types and positions of chemical groups in the molecular structure.^16^ Complex chemical structures can be digitized into a collection of molecular fingerprints (MFs) represented by binary bits, such as Molecular Access System (MACCS) keys.^17^ This information can serve as input for machine learning (ML) models to facilitate large-scale computational screening of drug compounds.^18^

Many drug-related chemical databases provide this input. For example, the Toxin and Toxin Target Database^19^ collects data from more than 18,000 sources, including other databases, government documents, books, and scientific literature. The toxins in this database are defined as chemicals that have been identified as hazardous in relatively low concentrations (<1 mM to <1 μM). They appear on multiple toxin and poison lists provided by government agencies or in the toxicological and medical literature. For each chemical, toxicity is evaluated by inspecting measurements such as minimum lethal dose or concentration and carcinogenicity in animal studies. Another database, ClinicalTrials.gov,^20^ collects information about clinical studies conducted around the world, including specific drugs under investigation, clinical trial status, study start and completion dates, and so on. The DrugBank^21^ database collects information from sources around the world, such as the FDA, Health Canada, European Medicines Agency, and ClinicalTrials.gov, to create a global knowledge base of drugs categorized as approved, withdrawn, or investigational.

Our goal in this study was to develop a filtering tool that can predict whether a small-molecule drug candidate will be worthy of clinical trials and whether it will be approved by the FDA. We collected information on a set of small-molecule compounds from the aforementioned drug-related databases and curated them into mutually exclusive categories based on their status in the drug discovery process. We then compared the distributions of 18 PCPs and a set of 166 MACCS MFs of these compounds and evaluated the ability of Lipinski’s Rule of Five, the Ghose filter, the Veber filter, the Rule of Three, and the REOS filter to identify unworthy compounds. Finally, we built classification models for these categories using a random forest ML model. The result is a computational tool that can predict FDA approvability of small-molecule drug compounds.

## RESULTS

### Curating Datasets of Toxic Compounds, Drugs in Clinical Trial, and FDA-approved Drugs

For every compound that survives the rigorous drug discovery process, thousands of candidates are eliminated. With the aim of developing a filtering tool to identify successful candidates and their associated PCPs, we collected information on thousands of small-molecule compounds from various drug-related databases. We then curated this information into three mutually exclusive groups for subsequent comparison and ML analyses: (1) the toxic compound (TD) group contained toxic compounds that would not be considered for clinical trial, as well as any preclinical research drugs that failed to enter clinical trial; (2) the clinical trial drug (CD) group included drugs that entered clinical trials, and (3) the FDA-approved (MD) group included drugs that were approved by the FDA, including those available on the market (MDon) or removed from the market (MDoff). In addition, we combined the CD and MD groups to form a group of non-toxic drugs (non-TD). To clarify, non-TD here refers to any drug eligible for clinical trials. The notation of non-TD serves simply to oppose TD; it does not imply that a drug is not toxic, since, for example, all drugs are toxic when overdosed. For convenience we may refer compounds in the TD group as toxic and in the non-TD group as non-toxic.

Toxicity information was collected from the Toxin and Toxin Target Database.^19^ Information on clinical and approved drugs was collected from DrugBank.^21^ Clinical trial status was obtained from ClinicalTrial.gov.^20^ All these data and information were retrieved before July 20, 2019. To ensure all selected compounds were small, we excluded biologics with a molecular weight exceeding 1,500 Daltons. We ensured that all selected compounds had profiles for the 18 PCPs analyzed. We also ensured that the fragments of chemical structures could be encoded with 166 bits of the MACCS keys.^17^ Our final dataset thus contained 2,567 compounds in the TD group, 1,463 in the CD group, and 2,133 (1,923 MDon and 210 MDoff) in the MD group.

### T-stochastic Neighbor Embedding Visualization of Toxic, Clinical Trial, and FDA-approved Compounds

We conducted a t-stochastic neighbor embedding (t-SNE) analysis to gauge the extent to which small-molecule drug compounds could be distinguished by their PCPs and MACCS MFs at different stages of the drug development process. t-SNE is a dimensionality reduction algorithm that uses matrices of pairwise similarities to visualize high-dimensional datasets.^22^ As shown in Figure 1, whereas compounds in the TD group were largely distinguishable from those in the non-TD group, the differences between drugs in the CD and MD groups or between MDon and MDoff were less obvious. These results indicate that accurate classification of the latter groups would be difficult to identify via simple rules or methods or even by sophisticated algorithms.

**Figure 1.**
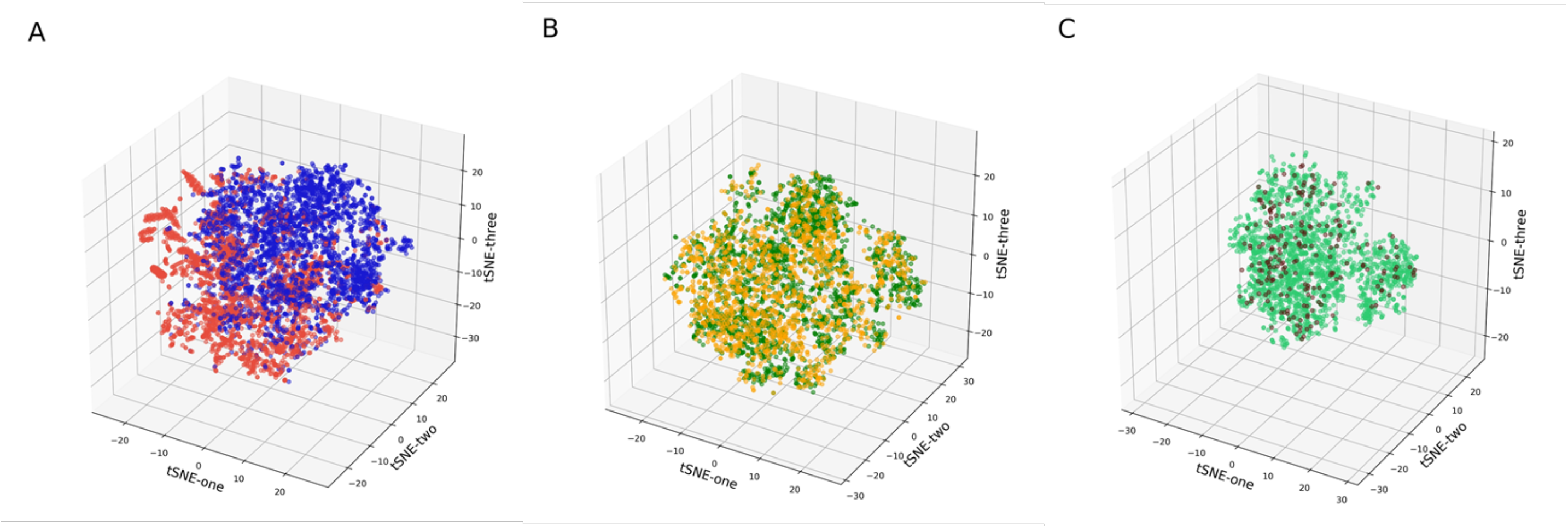
Three-dimensional t-stochastic neighbor embedding (t-SNE) analysis of toxic compounds, drugs in clinical trial, and FDA-approved approved drugs both on and off the market, using 184 features (18 physicochemical properties and 166 Molecular Access System molecular fingerprints). (A) non-toxic compounds (blue) versus toxic compounds (red); (B) FDA-approved drugs (green) versus drugs in clinical trials (orange); (C) drugs currently on the market (green) versus approved drugs withdrawn from the market (brown).

### Comparison of Physicochemical Properties

The PCPs of small-molecule compounds are important factors in drug design and development. To determine which of the 18 PCPs examined might be linked with successful drug development, we compared statistics for (1) toxic versus non-toxic drugs, (2) drugs in clinical trial versus FDA-approved drugs, and (3) approved drugs on the market versus those removed from the market. Figure 2 presents six examples of PCPs with significant differences in each of these three comparisons (see Supplementary Figure S1 for comparisons of the remaining PCPs). Some consistent trends were observed. For example, as expected, the overall PCP values for drugs in clinical trials, FDA-approved drugs, and non-toxic drugs were similar to each other but different from those of the toxic drugs. Intriguingly, compared to the PCPs of MDon drugs, the PCPs of MDoff drugs were more similar to those of the TD drugs, which might explain why they were withdrawn from the market (e.g., toxicity was not identified prior to FDA approval).

**Figure 2.**
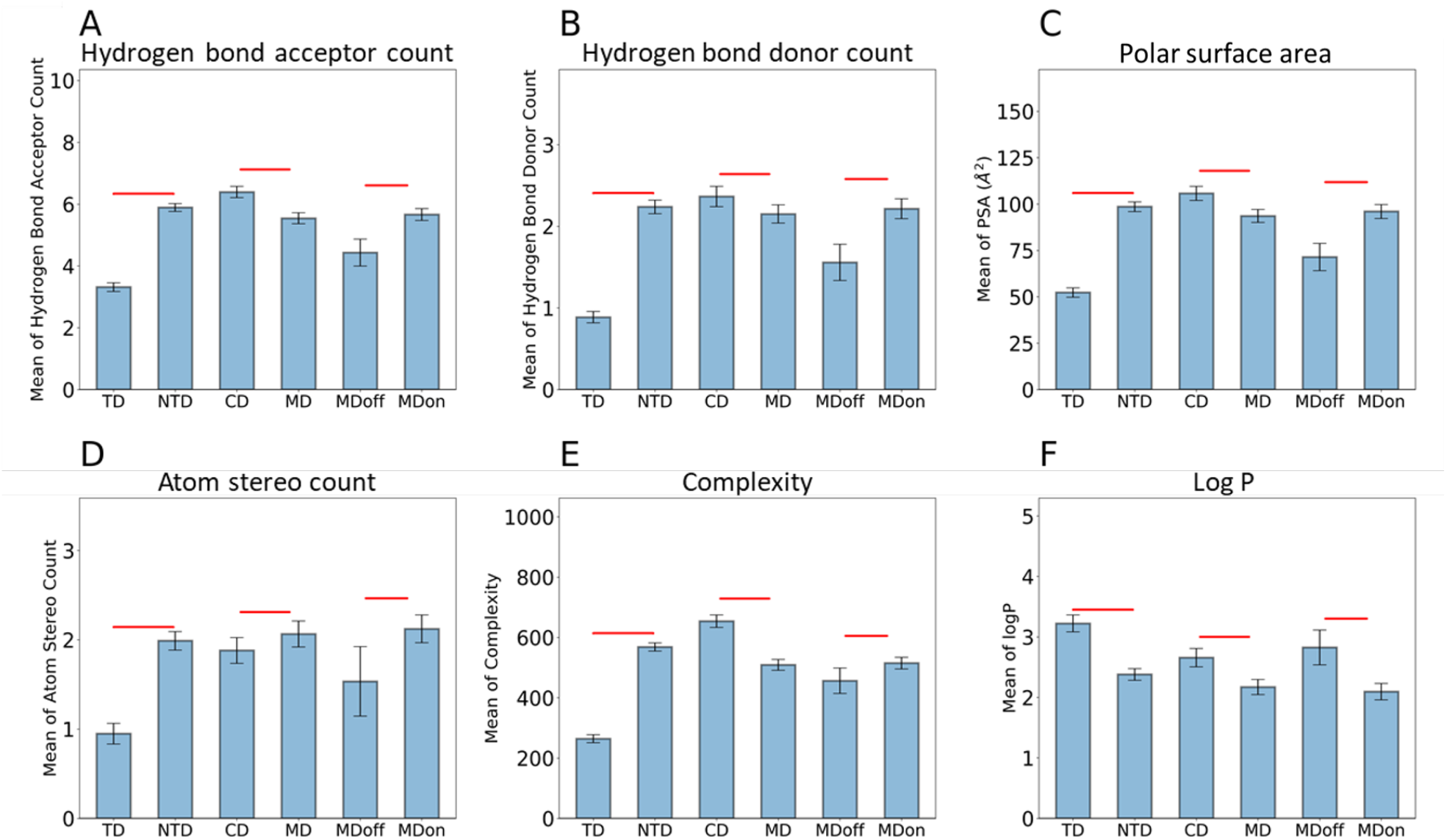
Physicochemical property comparisons of studied compounds. Bar plots show mean of each property in each compound group. Differences between two groups were tested using a one-tailed Student’s t-test. Error bar: 95% confidence interval. Red significance bridge: p value of t-test < 0.05. TD, toxic compound; NTD, non-toxic compound; CD, drug in clinical trial; MD, FDA-approved drug; MDoff, drug withdrawn from the market; MDon, drug currently on the market.

Some of our results help confirm findings reported in the literature. For example, we found that drugs classified as toxic and drugs taken off the market had significantly lower means of hydrogen bond acceptors and donors (Figure 2A and 2B), which may contribute to higher permeability and thus an increased possibility of adverse effects.^23^ In addition, a low polar surface area can result in drugs interacting with multiple receptors, which contributes to toxicity.^24^ In our study, the toxic compounds and the drugs no longer on the market had significantly lower polar surface area means than the other compounds. On the other hand, a large polar surface area can reduce drug efficacy and thus halt a clinical trial,^5^ and we observed that the mean polar surface areas of drugs in clinical trials were significantly higher than those for FDA-approved drugs (Figure 2C). We also observed high Log P values in the TD, CD, and MDoff groups (Figure 2F), which could be linked to adverse effects resulting from high plasma protein binding or tissue accumulation.^25^ The molecular weight of a drug compound usually increases during optimization. However, excessive molecular weight can inhibit solubility and permeability.^9^ Smaller drugs usually have better pharmacokinetic profiles.^26, 27^ Our analysis revealed that drugs in clinical trial had significantly higher molecular weight means than drugs that were FDA-approved (Supporting Information Figure S1J), perhaps explaining why many of their clinical trials were terminated.

### Comparison of Molecular Access System Molecular Fingerprints

We next analyzed the presence or absence of 166 MACCS MFs^17^ in these compounds (non-TD vs. TD, MD vs. CD, and MDon vs. MDoff). Table 1 lists the MFs for which the presence in one compound group was statistically significantly greater than in the other, based on a two-proportion Z-test. MFs are categorized as non-TD>TD, MD>CD, and MDon>MDoff in the left column and TD>non-TD, CD>MD, and MDoff>MDon in the right column, where > means the MF’s presence in the left-hand-side compound group is significantly greater. In this table, we exclude any MF exhibiting statistical significance in both columns of the comparisons. That is, Table 1 lists only the MFs that were consistently over-represented in compound groups that were favored (left column) or unfavored (right column) in the three stages that determine success or failure in the drug development process.

**Table 1.** Molecular Access System molecular fingerprints with significant differences between non-toxic, toxic, clinical trial, and FDA-approved (on and off the market) compounds.

| Comparison | key# | MACCS Fingerprint | Comparison | key# | MACCS Fingerprint |
| --- | --- | --- | --- | --- | --- |
| non-TD>TD | 8 | QAAA@1 | TD>non-TD | 3 | GROUP IVA, VA, VIA PERIODS 4-6 |
|  | 11 | 4M RING |  | 4 | ACTINIDE |
|  | 14 | S-S |  | 5 | GROUP IIIB,IVB |
|  | 17 | CTC |  | 16 | QAA@1 |
|  | 19 | 7M RING |  | 21 | C=C(Q)Q |
| | 26 | C\$=C(\$A)\$A | | 23 | NC(O)O |
|  | 32 | CSN |  | 31 | QX |
|  | 33 | NS |  | 34 | CH2=A |
|  | 36 | S HETEROCYCLE |  | 41 | CTN |
|  | 45 | C=CN |  | 46 | BR |
|  | 47 | SAN |  | 56 | ON(O)C |
|  | 50 | C=C(C)C |  | 63 | N=O |
|  | 59 | Snot%A%A |  | 103 | CL |
|  | 66 | CC(C)(C)A |  | **107** | **XA(A)A** |
|  | 74 | CH3ACH3 |  | **134** | **X (HALOGEN)** |
|  | 76 | C=C(A)A |  |  |  |
|  | 78 | C=N |  |  |  |
|  | 81 | SA(A)A |  |  |  |
|  | 84 | NH2 |  |  |  |
|  | 89 | OAAAO |  |  |  |
|  | 93 | QCH3 |  |  |  |
|  | 99 | C=C |  |  |  |
|  | 108 | CH3AAACH2A |  |  |  |
|  | 112 | AA(A)(A)A |  |  |  |
|  | 115 | CH3ACH2A |  |  |  |
|  | 116 | CH3AACH2A |  |  |  |
|  | 123 | OCO |  |  |  |
|  | **139** | **OH** |  |  |  |
| MD>CD | 8 | QAAA@1 | CD>MD | 4 | ACTINIDE |
|  | 11 | 4M RING |  | 21 | C=C(Q)Q |
| | 26 | C\$=C(\$A)\$A | | 23 | NC(O)O |
|  | 30 | CQ(C)(C)A |  | 24 | N-O |
|  | 39 | OS(O)O, 40 S-O |  | 41 | CTN |
|  | 45 | C=CN |  | 46 | BR |
|  | 50 | C=C(C)C |  | 71 | NO |
| | 76 | C=C(A)A | | 87 | X!A\$A |
|  | 115 | CH3ACH2A |  | **107** | **XA(A)A** |
|  | 116 | CH3AACH2A |  | 124 | QQ |
|  | 123 | OCO |  | **134** | **X (HALOGEN)** |
|  | **139** | **OH** |  |  |  |
| MDon>MDoff | 70 | QNQ | MDoff>MDon | 87 | X!A\$A |
|  | 89 | OAAAO |  | 103 | CL |
|  | **139** | **OH** |  | **107** | **XA(A)A** |
|  |  |  |  | **134** | **X (HALOGEN)** |

Table 1 shows that of the 166 MACCS MFs, only hydroxyl (OH, #139) and halogen (X, #107 and #134) exhibited significant differences in all three comparisons, with hydroxyl occurring more frequently in the favorable compounds list and halogen more frequently in unfavorable compounds list. The medicinal chemistry literature offers ample support for these findings. Adding a hydroxyl group to a chemical can shift metabolism from cytochrome P450 to conjugative enzymes, the products of which are not bioactive and are readily eliminated, leading to greatly increased polarity and thus less toxicity.^28^ One reason that halogenated structures occur more often in unfavorable compounds may be that carbon-halogen bonds are not easily metabolized, which could cause liver toxicity.^29^ Additionally, although it passed the statistical test for significance in only one of the three classification comparisons, compounds containing nitro (N-O, #24) and nitroso (N=O, #63) can undergo biological reduction to produce toxic metabolites that damage cells.^30, 31^ The toxicity of Groups IVA, VA, and VIA Periods 4-6 (4–6 row elements in the periodic table) (MACCS key #3) is well documented by their prevalence in toxic compounds.^32-36^

Boldface indicates favorable (left column) or unfavorable (right column) compounds in all three comparisons. X indicates halogen. % indicates aromatic query bond. A indicates any valid periodic table element symbol. Atoms bonded to the central atom are listed in parentheses. @ denotes a ring linkage, and the number following specifies the atom’s position in the line (e.g., @1 means linked back to the first atom in the list). ! denotes a chain or non-ring bond, and ! before a bond type specifies a chain bond. Q indicates heteroatoms (i.e., any non-C or non-H atom). $ indicates a ring bond. T denotes a triple bond, and not% means an aromatic query bond. M RING is the specific member of the ring, and # is the MACCS key number. MACCS, Molecular Access System; TD, toxic compound; CD, drug in clinical trial; MD, drug approved by FDA; MDoff, drug withdrawn from the market; MDon, drug currently on market.

### Performance of Simple Rules

PCP-based simple rules, such as the Rule of Five and others, have been widely employed to assist in designing and developing drug compounds.^9-13^ Here, we examined their ability to differentiate (1) non-toxic compounds from toxic, (2) FDA-approved (on and off the market) from clinical trial compounds (3) on market drugs from off market drugs. Table 2 shows that the five simple rules generally performed poorly in these classifications, even when comparing drugs in the TD and non-TD groups. Only the Ghose and REOS filters yielded respectable Matthews correlation coefficients (a performance measure for binary classification) of ∼0.3 for the comparisons. The Matthews correlation coefficients for the remaining comparisons were all lower than 0.2, and some were negative, meaning their discriminating power was worse than that for random guessing. These results suggest that adherence to well-known simple rules may not indicate a compound’s likelihood of FDA approval or its marketability. The results also cast doubt on simple rules’ ability to reliably screen compounds for preclinical toxicity tests.

**Table 2.** Sensitivity and specificity of physicochemical property-based simple rules and our machine learning model for differentiating between non-toxic, toxic, FDA-approved (on and off the market), and clinical trial compounds.

|  | non-TD vs. TD |  |  | MD vs. CD |  |  | MDon vs. MDoff |  |  |
| --- | --- | --- | --- | --- | --- | --- | --- | --- | --- |
|  | sen | spe | MCC | sen | spe | MCC | sen | spe | MCC |
| Rule of Five | 0.43 | 0.62 | 0.06 | 0.47 | 0.63 | 0.11 | 0.46 | 0.41 | -0.07 |
| Ghose filter | 0.40 | 0.90 | 0.33 | 0.37 | 0.55 | -0.08 | 0.36 | 0.58 | -0.03 |
| Veber filter | 0.76 | 0.10 | -0.17 | 0.77 | 0.25 | 0.03 | 0.76 | 0.11 | -0.09 |
| Rule of Three | 0.09 | 0.86 | -0.07 | 0.11 | 0.93 | 0.09 | 0.11 | 0.87 | -0.01 |
| REOS filter | 0.48 | 0.82 | 0.31 | 0.46 | 0.49 | -0.04 | 0.45 | 0.43 | -0.07 |
| Machine learning | 0.92 | 0.82 | 0.74 | 0.83 | 0.50 | 0.35 | 0.56 | 0.63 | 0.11 |
| Random guess | 0.5 | 0.5 | 0 | 0.5 | 0.5 | 0 | 0.5 | 0.5 | 0 |

The results of machine learning are the mean values of those from 10 cross-validation models; standard deviations are reported in the next section. sen: sensitivity; spe: specificity; MCC: Matthews correlation coefficient; TD, toxic compound; CD, drug in clinical trial; MD, drug approved by FDA; MDoff, drug withdrawn from the market; MDon, drug currently on market; REOS, Rapid Elimination of Swill.

### Results of Random Forest Machine Learning

To investigate whether a sophisticated classification algorithm could significantly outperform these simple rules, we conducted 10-fold cross-validation ML using the random forest algorithm.^37^ Using 184 dimension features (18 PCPs + 166 MACCS MFs) for each compound, we derived one ML model for each of the three classifications (see Materials and Methods). Figure 3 shows their performances on test sets, revealing that the ML models performed best at classifying non-toxic versus toxic compounds, followed by FDA-approved versus clinical trial compounds, and they produced marginally significant results for differentiating between FDA-approved drugs on versus off the market. The areas under the curve of the receiver operating characteristics across the 10 independent ML models for the three classifications were respectively 0.94 ± 0.01 (mean ± standard deviation), 0.73 ± 0.02, and 0.62 ± 0.06; the sensitivities were 0.92 ± 0.01, 0.83 ± 0.01, and 0.56 ± 0.06; the specificities were 0.82 ± 0.02, 0.50 ± 0.04, and 0.63 ± 0.13; and the precisions were 0.88 ± 0.01, 0.71 ± 0.02, and 0.93 ± 0.02. Table 2 presents the respective MCCs and their means (0.74 ± 0.02, 0.35 ± 0.04, and 0.11 ± 0.08). The precision results of the ML models indicate an average compound attrition rate of 12% for preclinical compounds and 29% for compounds in clinical trials. The variations, or model uncertainties, in these test predictions increased moderately from the first classification to the second and greatly from the second to the third. These results highlight the increasing difficulty in filtering out unfavorable compounds in the drug development process. In comparison, simple rules performed much worse and more erratically than our ML model did in classifying drug compounds.

**Figure 3.**
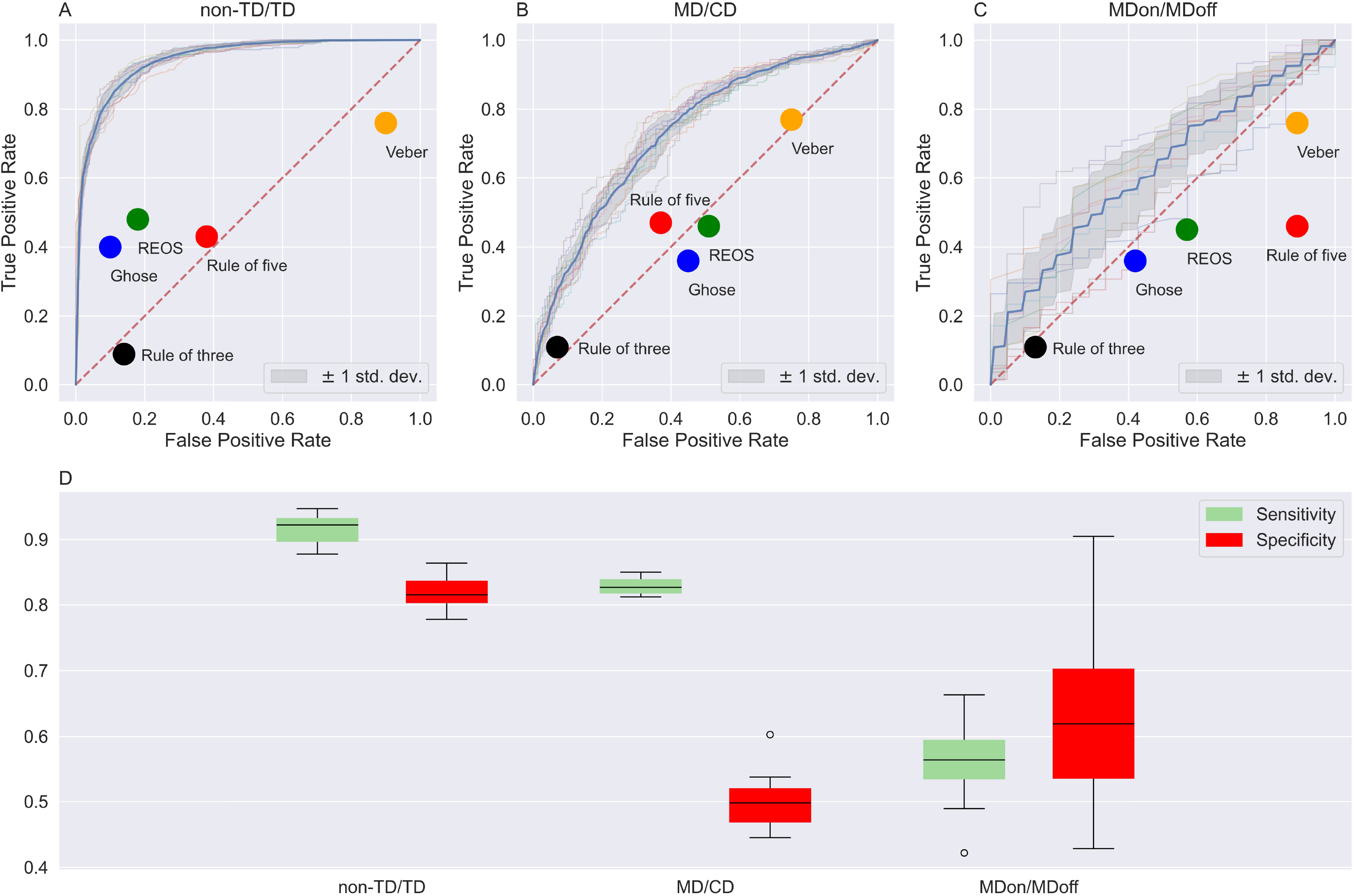
Model performance, as derived from 10-fold cross-validation random forest machine learning. The models’ prediction results are based on test set compounds for the three classification comparisons (non-TD/TD, MD/CD, and MDon/MDoff). The top panels show the receiver operating characteristic curves (gray line) for 10 independent models and their mean receiver operating characteristic (blue line) on the classification of (A) non-TD/TD, (B) MD/CD, and (C) MDon/MDoff. Color dots indicate the performance of various simple rules. (D) Boxplots of model sensitivity and specificity, computed using a 0.5 probability threshold for the three classification comparisons. The true positive rate is sensitivity, and the false positive rate is 1-specificity. TD, toxic compound; CD, drug in clinical trial; MD, drug approved by FDA; MDoff, drug withdrawn from the market; MDon, drug currently on the market.

We further investigated the predictive power of models derived using only the 18 PCPs or only the 166 MACCS MFs as model features. The results, shown in Supplementary Figure S3 and Figure S4, indicated that PCPs were much more predictive than MFs for the three classification tasks investigated, but the combination of both was required to achieve the full prediction benefit, as shown in Figure 3. We confirmed this observation by analyzing feature importance in the random forest algorithm, which calculated the average decrease in impurity across trees.^38^ As illustrated in Figure 4, most of the 18 PCPs contributed to the model’s predictions with recognizable significance for at least one of the three classification comparisons, but very few MFs contributed similarly. Results from the feature importance analysis generally aligned with findings from the previously described statistical analysis. For example, Log P, polar surface area, complexity, and numbers of hydrogen bond donors and acceptors were prominent, and the presence of halogen helped discriminate between favorable and unfavorable compounds. The analysis also revealed differential importance of certain features at different stages of the drug development process. Understanding the significance of these results requires further investigation. We further tested our ML models using more recent information from DrugBank posted on

**Figure 4.**
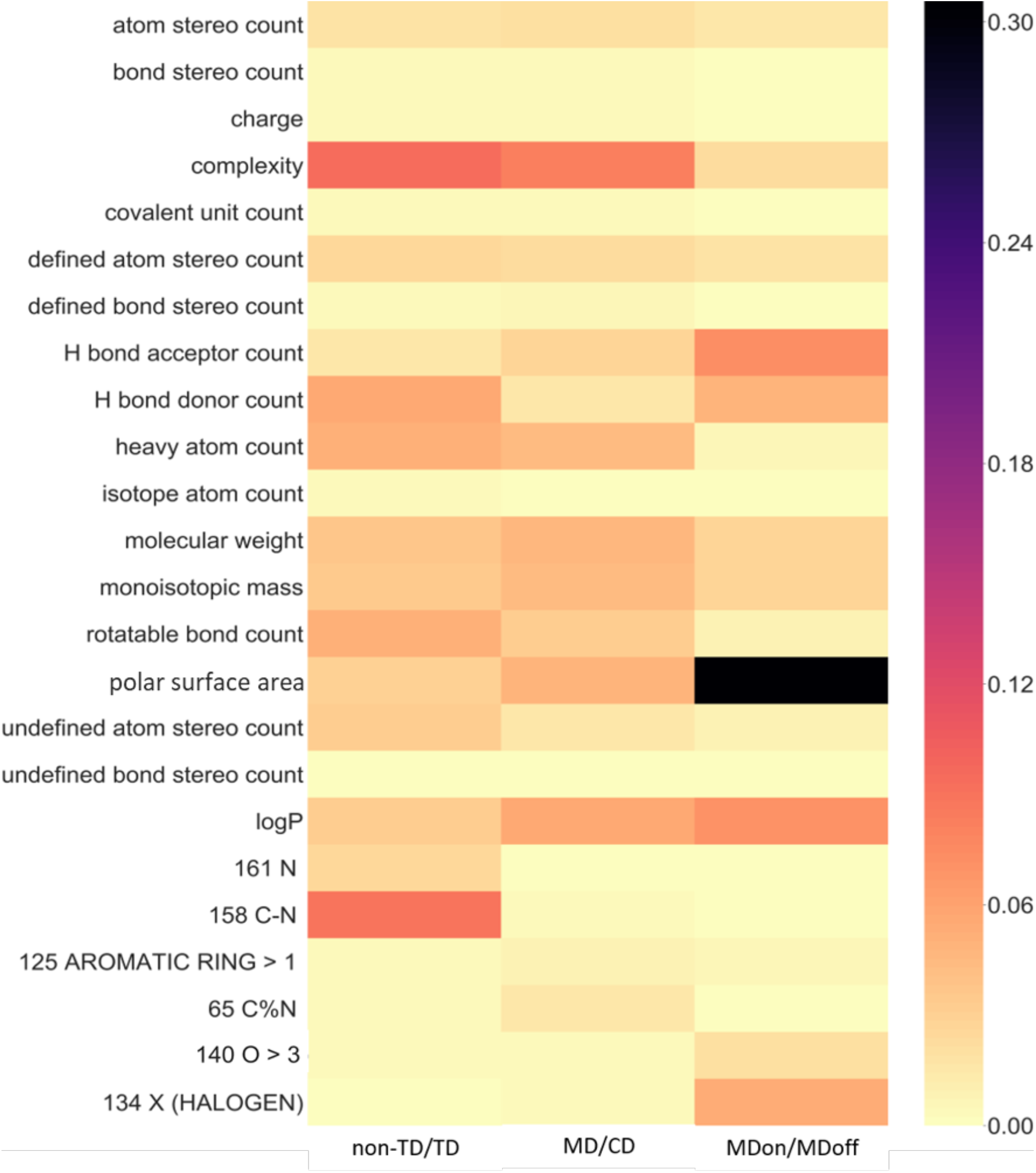
Heat map of feature importance for the machine learning models to discriminate compounds in each of the three classification comparisons. To the right is the importance score bar (darker colors indicate more importance). See Durant et al.’s report^17^ and the RDkit^39^ GitHub repository for the identification number and definition of the Molecular Access System molecular fingerprints.

July 2, 2020, which is nearly one year after the date we originally downloaded the database. We found five new FDA-approved drugs, six new drugs in clinical trials, and 27 drugs from clinical trials that had been subsequently approved by the FDA, all with molecular weights below 1,500 Daltons. We predicted the compound class for each drug that was newly FDA-approved or in clinical trial using the 10 cross-validation models and choosing their consensus as the predicted class. The 27 newly approved compounds were still in clinical trial in the database that we downloaded to develop the ML models for classifying non-toxic/toxic and FDA-approved/clinical trial drugs. These compounds were present in the test fold for only one of the 10 models and could be predicted only by that particular model (see Materials and Methods). However, to classify drugs as currently on the market or off the market, we could treat them as new compounds to be predicted by all 10 MDon/MDoff models. Information on clinical trials for the five newly FDA-approved drugs was not in the original database used to train the ML models; these drugs were treated as new compounds for all three classification predictions.

As the results in Table 3 show, our non-TD/TD models predicted all five newly FDA-approved drugs and all six drugs in clinical trials to be eligible for clinical trials. These findings validate the reliability of our ML models in predicting compound eligibility for a clinical trial. As expected, the ability of the ML models to predict FDA approval among drugs in clinical trials diminished significantly, identifying four out of five (80%) newly approved drugs (Table 3) and predicted 11 out of the 27 drugs (41%) in clinical trials to be approved (Table 4). Nevertheless, these prediction rates are much higher than the 10%–20% FDA approval rate among compounds in clinical trials. Similar results were obtained for drugs not collected in our compound datasets but were approved for use in Europe or in Japan, with the non-TD/TD models producing a sensitivity of 0.96 (246/256) and the MD/CD models a sensitivity of 0.62 (158/256) for those approved drugs (Supplemental Table S4).

**Table 3.** Machine learning models’ ability to predict newly FDA-approved drugs and new compounds eligible for clinical trial.

|  | Compound name | PubChem ID | non-TD/TD | MD/CD | MDon/MDoff |
| --- | --- | --- | --- | --- | --- |
| New<br>MDs | Levamlodipine | 9822750 | non-TD (10) | MD (10) | MDon (7) |
|  | Prednisolone acetate | 5834 | non-TD (10) | MD (10) | MDon (10) |
|  | Taurocholic acid | 6675 | non-TD (10) | MD (7) | MDon (10) |
|  | Metergoline | 28693 | non-TD (10) | CD (10) | MDoff (10) |
|  | Xamoterol | 155774 | non-TD (10) | MD (10) | MDon (9) |
| New<br>CDs | RO-4584820 | 6918852 | non-TD (10) | CD (10) | MDon (9) |
|  | Prinaberel | 5388909 | non-TD (10) | MD (10) | MDoff (10) |
|  | Lidorestat | 157839 | non-TD (10) | CD (10) | MDoff (10) |
|  | PF-2545920 | 11581936 | non-TD (10) | CD (10) | MDoff (9) |
|  | SD-06 | 9865587 | non-TD (10) | CD (10) | MDon (8) |
|  | 622402-22-6 | 9956726 | non-TD (10) | CD (10) | MDon (6) |

Number in parenthesis indicates the number of models predicting the class. TD, toxic compound; CD, drug in clinical trial; MD, drug approved by FDA; MDoff, drug withdrawn from the market; MDon, drug currently on the market.

**Table 4.**
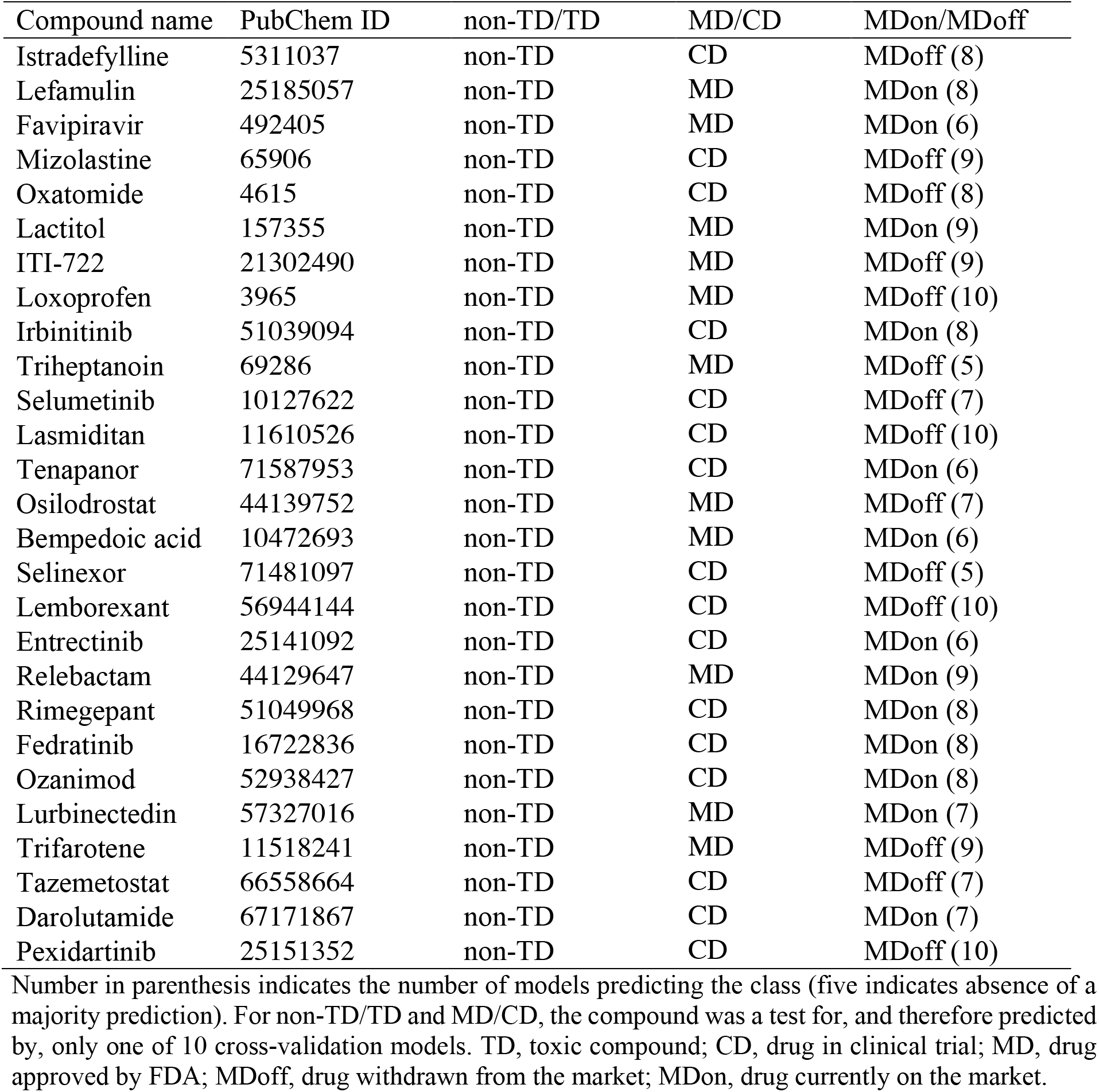
Machine learning models’ ability to predict compounds moving from clinical trial to FDA-approved, based on updated DrugBank data.

## DISCUSSION

The field of medicinal chemistry has accumulated considerable knowledge and physicochemical rules about drug compounds. Yet, most of these compounds do not pass preclinical and clinical examinations. High compound attrition rates contribute to a costly trial-and-error drug discovery process. In theory, a drug’s safety and efficacy are largely encoded in its molecular composition and structure, but the longstanding problem of compound attrition in drug discovery underscores the difficulty in modeling the complex relationships between compound structure and activity, which are further complicated by factors such as the drug target and potential interactions.

ML methods can effectively map these convoluted relationships to make accurate classifications and outcome predictions, albeit with difficulty explaining or describing the relationships. We found that the random forest ML method reduced compound attrition rates better than physicochemical rules by several folds (Table 2 and Figure 3). Encouragingly, our analysis of feature importance (Figure 4) revealed several molecular insights in accordance with the literature, although it remains difficult to deduce generally applicable and explicit rules. A similar but different ML drug classifier was proposed to predict likelihood of drugs showing toxicity in clinical trials.^40^ Comparing to our MD/CD models, its positive set also consisted of FDA-approved drugs, but only 100 drugs that failed in clinical trials for toxicity reason were used in the negative set. In addition, its prediction requires input of not only a drug’s physicochemical properties but also properties such as tissue expression of its target genes.^40^

The ML models’ ability to filter out ineligible compounds decreased over the course of the drug discovery process (Figure 3). Besides the decreasing number of compounds available for training the successive ML models, one reason for this decreased predictive ability was the use of the compound status annotated in the database to produce mutually exclusive compound groups for classification training. In reality, status annotations are not permanent. Thus, a portion of compounds in clinical trial that would be approved by the FDA and put on the market remained classified as clinical drugs to train the MD/CD models. A drug on the market also could re-enter clinical trial for a different disease, or the FDA could reapprove a drug no longer on the market for a particular patient cohort (e.g., specific genetic background). Such status changes would not significantly affect model performance, however, because, as the high attrition rates indicate, these cases represent only a small portion among a large number of failed compounds.

Another factor that likely contributed to the progressive deficiency of the model relates to inconsistent predictions among the three classification models. Consider the new compound eligible for clinical trial, RO-4584820 (Table 3). 10 out of 10 MD/CD models favored it to be a clinical trial drug over an FDA-approved drug, but 9 out of 10 MDon/MDoff models predicted it to stay on the market. Although we observed similar examples of inconsistent predictions, we also observed many consistent predictions. Among newly approved drugs and new drugs in clinical trial, many compounds (e.g. Levamlodipine and Prednisolone acetate) predicted by the MDon/MDoff model to succeed (i.e., MDon) were classified at an earlier stage with a consistent prediction (i.e., MD by the MD/CD models and non-TD by the non-TD/TD models).

Despite the high compound attrition rate of drug discovery, there isn’t yet a database available in the public domain that collects compounds deemed unsuitable for clinical trials whether or not they failed toxicity test in preclinical research. In this study, we used Toxin and Toxin Target Database to find clinically ineligible compounds with the justification that toxicity is the major reason for compounds to be excluded from clinical trials.^41^

These deficiencies and uncertainties notwithstanding, our ML models and the molecular insights revealed could provide useful guidelines to help medicinal chemists design, modify, or screen drug compounds. As drug-related databases expand to increase the number, chemical diversity, and status annotations of drug compounds, ML models are bound to improve prediction powers and become indispensable for drug discovery.

## CONCLUSION

We developed ML models to computationally predict which small-molecule compounds would be eligible for clinical trials and the likelihood that they would be approved by the FDA. Tests on compounds not used to train the models showed that viable clinical drugs and FDA approvability can be predicted a priori based solely on chemical structure and physicochemical properties, leading to a drastic reduction in the time-consuming and expensive attrition rate associated with drug discovery. If substantiated by further investigations, this approach could offer a practical and cost-effective method for use in the medicine, biotechnology, and pharmacology fields.

## MATERIALS AND METHODS

### Collections of Compounds and Molecular Features

From the Toxin and Toxin Target Database,^19^ we collected the PubChem IDs of 3,678 toxic compounds. PubChem IDs for 3,581 drugs currently on the market, 253 drugs off the market, and 2,497 drugs in clinical trial were retrieved from DrugBank.^21^ Using PubChemPy, a Python package, we collected the numerical values of 18 PCPs of these compounds from PubChem (charge, molecular weight, Log P, monoisotopic mass, polar surface area, complexity, H bond donor count, H bond acceptor count, rotatable bond count, heavy atom count, isotope atom count, atom stereocenter count, defined atom stereocenter count, undefined atom stereocenter count, bond stereocenter count, defined bond stereocenter count, undefined bond stereocenter count, and covalently-bonded unit count).^42^ Complexity (i.e., the elemental and structural features of the compound, such as symmetry) was computed using the Bertz/Hendrickson/Ihlenfeldt formula.^43^

In this report, we limited our analysis to small molecules as opposed to biologics. We removed compounds with molecular weights exceeding 1,500 Daltons based on the distribution of compound molecular weight in DrugBank. We thus obtained four mutually exclusive lists of compounds, including profiles of their 18 PCPs: (1) 2,567 toxic compounds (TD), (2) 1,463 drugs in clinical trial (CD), (3) 1,923 drugs on the market (MDon), and (4) 210 drugs no longer on the market (MDoff). To ensure mutual exclusiveness, when a compound was found present in multiple drug-related databases having different status annotations, we resolved the conflict by setting the priority MDoff > MDon > CD > TD.

The missing Log P values for 1,125 compounds were imputed using the mean value of Log P of all the other compounds in the set (training, validation, or testing) when building the ML models. The 166 bits of MACCS keys representing various MFs^17^ for all compounds were obtained via RDKit, a Python chemo-informatics package. Figure 5 shows a flowchart of our curation of the compound datasets. Full descriptions of the 166 MACCS MFs can be found in Durant et al.^17^ and RDkit’s GitHub repository.^39^

**Figure 5.**
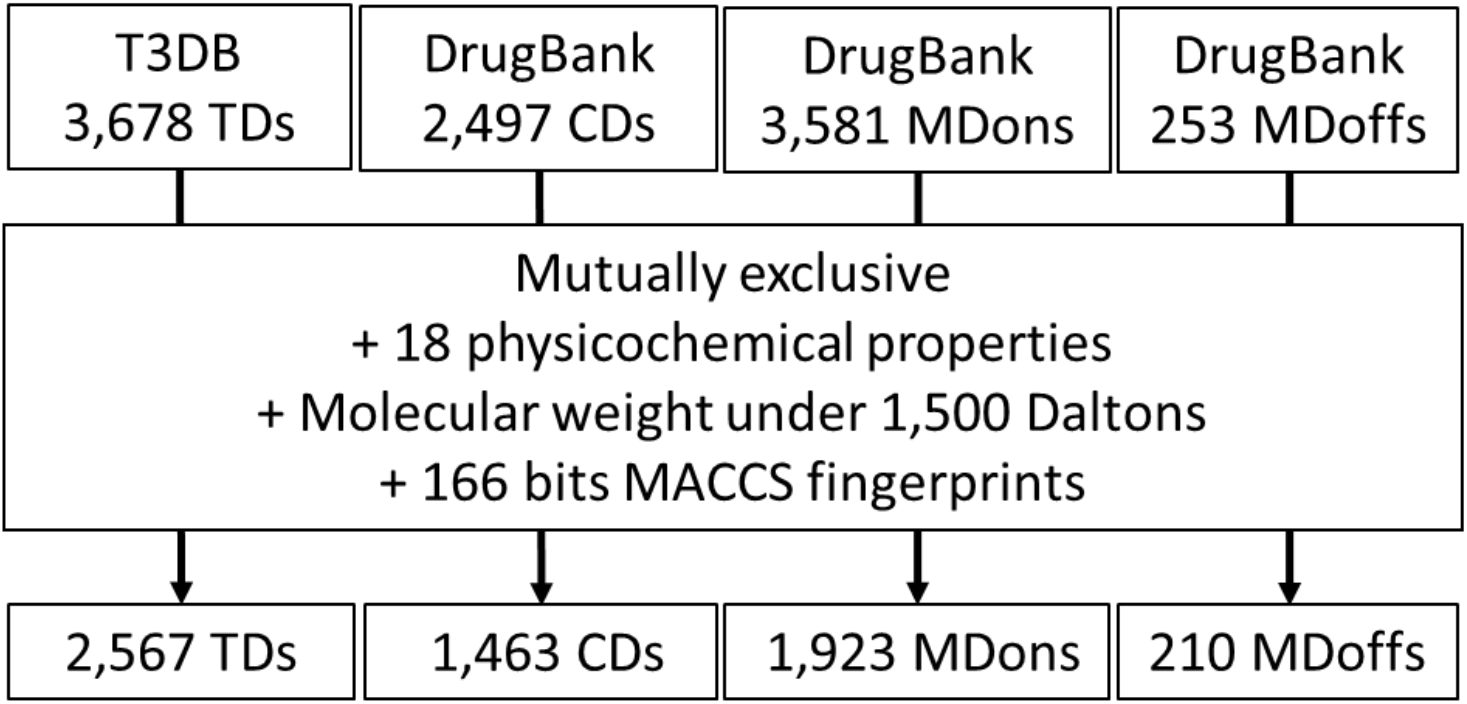
Flowchart of building exclusive datasets of small-molecule drug compounds. T3DB, Toxin and Toxin Target Database; TD, toxin; CD, in clinical trial; MDon, currently on the market; MDoff, withdrawn from the market; MACCS, Molecular Access System.

### Analysis of Physicochemical Properties and Molecular Access System Fingerprints of Drugs Identified as Toxins, in Clinical Trial, on the Market, and no Longer on the Market

To compare PCPs, the mean value of each PCP for each compound group was calculated. One-tailed student’s *t* tests were performed to test whether any PCP differed significantly between two compared compound groups, which were (1) toxic versus non-toxic drugs, (2) drugs in clinical trial versus FDA-approved drugs, and (3) approved drugs on the market versus those removed from the market. Non-toxic drugs (non-TD) are comprised of drugs in clinical trial and FDA-approved drugs. To compare MFs in different compound groups, we defined M_i_ as the proportion of compounds in a given compound group with the MACCS key (i.e., MF) i, the presence of which was represented by setting bit i = 1. The proportion was calculated using the equation M_i_ = C_i_/C_t_, where C_i_ is the number of compounds possessing MF i, and C_t_ is the total number of compounds in the compound group. Differences in proportions for each of the 166 MACCS MFs^17^ in the comparisons (non-TD vs. TD, MD vs. CD, and MDon vs. MDoff) were tested using a one-tailed two-proportion Z-test for statistical significance using the Python StatsModels package.^44^ P values less than 0.05 were regarded as statistically significant.

### Filtering Drug Compounds Using Simple Rules Based on Physicochemical Properties

Compounds meeting the Rule of Five,^9^ Ghose,^10^ Veber,^11^ Rule of Three,^12^ and REOS^13^ were determined according to their respective criteria, shown in Supplementary Table S1. To evaluate the power of these simple rules in differentiating between groups (non-TD vs. TD, MD vs. CD, and MDon vs. MDoff), we defined the non-TD, MD, and MDon groups as positive and the TD, CD, and MDoff group as negative in their respective comparisons. Sensitivity, specificity, MCC, and precision were calculated as follows. Note that by predicting the proportion of failed compounds (i.e., those falsely predicted to be in the positive group), 1-precision is a proxy for the compound attrition rate.

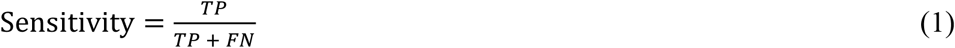

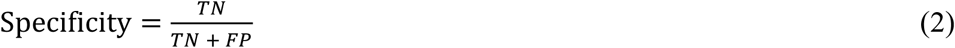

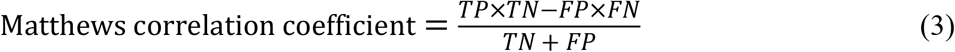

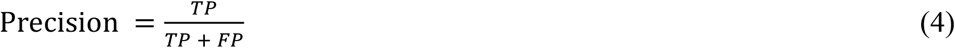

*where TP* = true positive, *TN* = true negative, *FP* = false positive, and *FN* = false negative.

### Building and Evaluating Machine Learning Models

Similarly, in deriving the three classification models of non-TD/TD, MD/CD, and MDon/MDoff, we defined non-TD, MD, and MDon as positive groups and TD, CD and MDoff as negative groups respectively. We used the random forest classifier in Scikit-learn,^45^ an ML library for Python, for this procedure. The compounds curated from drug-related databases, as shown in Figure 5, were randomly divided into 10 equal parts to establish an 8-1-1 training, validation, and test scheme of ML. For example, for the non-TD/TD classification, we included 2,877 non-toxic compounds and 2,053 toxic compounds in the training set, 359 non-toxic compounds and 257 toxic compounds in the validation set, and 360 non-toxic compounds and 257 toxic compounds in the test set. We balanced the training set and validation set to derive the MDon/MDoff model, because the number of compounds in the MDon group (1,923) was much greater than that in the MDoff group (210). Supplementary Table S2 lists the numbers of compounds used for training, validation, and testing to derive the ML models for the three classifications.

Model parameters were optimized using an internal, 10-fold, cross-validated random search over a parameter grid, shown in Supplementary Table S3. Models were evaluated on their performance in the validation set, and the best-performing model was chosen for the test set. Sensitivity was used to select the best-performing model for the non-TD/TD and MD/CD predictions. Specificity was used for the MDon/MDoff prediction, because high specificity can more accurately determine whether a drug will be pulled from the market. We repeated the procedure 10 times to complete cross-validation learning and testing. A rolling scheme was applied to the 10 equally divided data parts upon each repetition. This rolling data-segment scheme ensured that every compound occurred only once in the validation and only once in the testing among the 10-fold cross-validation models, thus avoiding data contamination.^46^ Performance of each of the resulting 10 models for each classification was evaluated based on their respective test set according to sensitivity (equation 1), specificity (equation 2), MCC (equation 3), and precision (equation 4), as well as the area under the receiver operating characteristic curve. We conducted a feature importance analysis by computing gini impurity to measure how much a feature was used by the tree nodes in the random forest model, thus indicating the extent of the feature’s importance to the model.^38, 47^

## DATA AVAILABILITY

All data are included in the manuscript and supporting information. Source code for the model is available at GitHub, https://github.com/ChihHanHuang/Predicting-FDA-approvability

## AUTHOR CONTRIBUTIONS

C.H.H. and M.J.H initiated and designed the study. C.H.H., J.H. and L.Y.Y. curated the datasets and performed statistical analysis. C.H.H. and J.H. developed ML models and analysis. T.M.C.,

E.S.C.S. and M.J.H supervised the study. C.H.H., J.H., L.Y.Y. and M.J.H wrote and edited the manuscript.

## ACKNOWLEDGMENTS

The authors thank Dr Kuo-Kan Liang and Dr. Szu-Hua Pan for their feedback at the early stage of the study; Dr. Wen-Shan Li, Dr. Jung-Hsin Lin and Dr. Sherry Ku for their suggestions; and past and current members of the Hwang lab for their discussions.

## ABBREVIATIONS

FDA: Food and Drug Administration
MACCS: Molecular Access System
MF: molecular fingerprint
ML: machine learning
PCP: physicochemical property
REOS: Rapid Elimination of Swill
t-SNE: t-stochastic neighbor embedding
TD: toxic compound
CD: drug in clinical trial
MD: drug approved by FDA
MDoff: drug withdrawn from the market
MDon: drug currently on the market

