## Supplementary tables and figures for "Predicting FDA approvability of small-molecule drugs"

**Contents**

Table S1. Criteria of simple rules based on physicochemical properties

Table S2. Numbers of compounds in the 10-fold, cross-validation to derive machine learning models

Table S3. Grid used for parameter tuning by the random forest machine learning

Table S4. Prediction results of Europe-approved drugs and Japan-approved Drugs

Figure S1: Physicochemical property comparison results between (1) TD vs. non-TD, (2) CD vs. MD, and (3) MDoff vs. MDon

Figure S2. Heat map of feature importance for all features used by the machine learning models

Figure S3. Model performance using only the 18 physicochemical properties

Figure S4. Model performance using only the 166 Molecular Access System fingerprints

**Table S1. Criteria of simple rules based on physicochemical properties**

| Filter | Criteria |
|---|---|
| Rule of Five[1] | Molecular weight < 500 Daltons<br>H-bond donor count ≤ 5<br>H-bond acceptor count ≤ 10<br>Log P ≤ 5 |
| Ghose filter[2] | Log P: -0.4 ~ 5.6<br>Molecular weight: 160 ~ 480 Daltons<br>Molar refractivity: 40 ~ 130<br>Total number of atoms: 20 ~ 70 |
| Veber filter[3] | Rotatable bonds ≤ 10<br>Topological polar surface area ≤ 140 $Å^2$ |
| Rule of Three[4] | Molecular weight ≤ 300 Daltons<br>Log P ≤ 3<br>H-bond donor count ≤ 3<br>H-bond acceptor count ≤ 3<br>Rotatable bonds ≤ 3 |
| Rapid Elimination of Swill (REOS) filter[5] | Molecular weight: 200 ~ 500 Daltons<br>Log P: -5.0 ~ +5.0<br>H-bond donor count: 0 ~ 5<br>H-bond acceptor count: 0 ~ 10<br>Formal charge: -2 ~ +2<br>Rotatable bond count: 0 ~ 8<br>Heavy atom count: 15 ~ 50 |

**Table S2. Numbers of compounds in the 10-fold, cross-validation to derive machine learning models**

| | non-TD/TD | MD/CD | MDon/MDoff |
|---|---|---|---|
| Training set | 2877 ± 1/2054 ± 1 | 1706 ± 1/1170 ± 1 | 168/168 |
| Validation set | 359+ 1/257 | 213 + 1/146 + 1 | 21/21 |
| Test set | 359 + 1/257 | 213 + 1/146 + 1 | 192 + 1/21 |

Number of compounds in each compound group with plus or minus indicating variation among folds. TD, toxic compound; CD, drug in clinical trial; MD, drug approved by FDA; MDon, drug currently on the market; MDoff, drug withdrawn from the market.

**Table S3. Grid used for parameter tuning by the random forest machine learning**

| Parameter | Grid value | Description |
|---|---|---|
| n_estimators | [100,500,1000] | Number of trees generated |
| max_features | [0.4, 0.5, 0.6] | Fraction of features to consider for the best split |
| min_samples_leaf | [1,50,100] | Minimum number of samples required at a leaf node |

**Table S4. Prediction results of Europe-approved drugs and Japan-approved Drugs**

| Drug name | PubChemID | Approval | non-TD/TD | MD/CD | MDon/MDoff |
|---|---|---|---|---|---|
| Corlentor | 3045381 | EU and JPN | non-TD (10) | CD (10) | MDoff (6) |
| Macugen | 56603655 | EU and JPN | non-TD (10) | CD (9) | MDon (10) |
| Eslax | 441351 | JPN | non-TD (10) | MD (10) | MDon (7) |
| Senstend | 9911821 | EU and JPN | non-TD (10) | MD (9) | MDon (5) |
| Spelear | 134669 | JPN | non-TD (10) | MD (10) | MDon (10) |
| Glufast | 5478927 | JPN | non-TD (10) | MD (10) | MDon (9) |
| Tasermity | 159247 | EU and JPN | non-TD (10) | MD (10) | MDoff (9) |
| Zafatek | 44183569 | JPN | non-TD (10) | CD (10) | MDon (9) |
| Fostoin | 56338 | JPN | non-TD (10) | CD (9) | MDon (9) |
| Ferinject | 86278165 | JPN | non-TD (10) | MD (10) | MDon (10) |
| Champix | 6918678 | EU and JPN | non-TD (10) | MD (10) | MDon (10) |
| Pemazyre | 86705695 | EU and JPN | non-TD (10) | CD (10) | MDon (8) |
| Ameparomo | 441375 | JPN | non-TD (10) | MD (10) | MDon (10) |
| Jzoloft | 63009 | JPN | non-TD (10) | MD (10) | MDoff (8) |
| Normosang | 135564839 | JPN | non-TD (10) | MD (5) | MDon (10) |
| Xeljanz | 10174505 | EU and JPN | non-TD (10) | CD (7) | MDon (10) |
| Lynparza | 23725625 | EU and JPN | non-TD (10) | CD (10) | MDoff (7) |
| Azilva | 135415867 | JPN | non-TD (10) | CD (10) | MDon (7) |
| Urece | 51349053 | JPN | non-TD (10) | MD (10) | MDoff (10) |
| Ceredist | 114750 | JPN | non-TD (10) | CD (8) | MDon (10) |
| Trintellix | 56843850 | JPN | non-TD (10) | MD (10) | MDoff (8) |
| Ongentys | 135565903 | EU and JPN | non-TD (10) | CD (10) | MDon (6) |
| Deltyba | 6480466 | EU and JPN | non-TD (10) | CD (10) | MDon (8) |
| Gracevit | 461399 | JPN | non-TD (10) | CD (7) | MDoff (6) |
| PASIL, | 6918232 | JPN | non-TD (10) | CD (8) | MDon (7) |
| Smyraf | 67998300 | JPN | non-TD (10) | CD (10) | MDon (10) |
| Rolufta Ellipta | 11519069 | EU and JPN | non-TD (10) | MD (10) | MDon (5) |
| Opatanol | 5282402 | EU and JPN | non-TD (10) | MD (10) | MDoff (8) |
| Velexbru | 71571562 | JPN | non-TD (10) | CD (10) | MDon (8) |
| Musredo | 72710763 | JPN | non-TD (10) | MD (6) | MDon (9) |
| Nailin | 154723947 | JPN | non-TD (10) | MD (6) | MDon (10) |
| Cerdelga | 52918379 | JPN | non-TD (10) | MD (9) | MDon (10) |
| Votrient | 11525740 | EU and JPN | non-TD (10) | CD (10) | MDon (10) |
| Anerem | 23658607 | JPN | non-TD (10) | CD (10) | MDon (10) |
| Riamet | 6450800 | JPN | non-TD (9) | MD (10) | MDon (8) |
| Entecavir Accord | 135398508 | EU and JPN | non-TD (10) | MD (5) | MDon (10) |
| Livostin | 54384 | JPN | non-TD (10) | MD (10) | MDoff (6) |
| Giotrif | 15606394 | EU and JPN | non-TD (10) | CD (9) | MDon (10) |
| Ecclock | 86301316 | JPN | non-TD (10) | MD (10) | MDon (7) |
| nan | 445063 | EU and JPN | non-TD (10) | MD (10) | MDon (10) |
| Defitelio | 135565962 | EU and JPN | non-TD (10) | CD (10) | MDon (10) |
| Zerbaxa | 86291594 | EU and JPN | non-TD (10) | MD (10) | MDon (9) |
| Laventair | 71300746 | EU and JPN | non-TD (10) | MD (10) | MDon (10) |
| Pheburane | 5258 | EU and JPN | TD (10) | MD (10) | MDoff (6) |
| Lexapro | 146571 | JPN | non-TD (10) | MD (9) | MDon (10) |
| Nopicor | 6918287 | JPN | non-TD (10) | CD (9) | MDon (10) |
| Viviant | 154256 | JPN | non-TD (10) | CD (10) | MDon (10) |
| GHRP | 6321294 | JPN | non-TD (10) | CD (10) | MDon (9) |
| Vegamox | 101526 | JPN | non-TD (10) | CD (8) | MDon (7) |
| Crestor | 5282455 | JPN | non-TD (10) | MD (6) | MDon (9) |
| Abiraterone Accord | 9821849 | EU and JPN | non-TD (10) | MD (10) | MDoff (6) |
| Copaxone | 3081884 | JPN | non-TD (10) | MD (10) | MDon (10) |

| | | | | | |
|---|---|---|---|---|---|
| Harvoni | 72734365 | EU and JPN | non-TD (10) | MD (8) | MDon (8) |
| Lullan | 115368 | JPN | non-TD (10) | CD (10) | MDoff (6) |
| Talion | 164521 | JPN | non-TD (10) | MD (7) | MDon (10) |
| Vyndaqel | 24970412 | JPN | non-TD (10) | CD (10) | MDon (10) |
| Maxalt | 77997 | JPN | non-TD (10) | CD (8) | MDon (7) |
| Tamiflu | 78000 | EU and JPN | non-TD (10) | MD (10) | MDon (10) |
| Tepmetko | 25171648 | JPN | non-TD (10) | CD (10) | MDon (6) |
| Aiphagan | 6917831 | JPN | non-TD (10) | MD (5) | MDon (9) |
| Jakavi | 25127112 | EU and JPN | non-TD (10) | CD (10) | MDon (10) |
| Calsed | 114897 | JPN | non-TD (10) | MD (10) | MDon (10) |
| Memantine Accord | 181458 | EU and JPN | non-TD (10) | MD (10) | MDoff (8) |
| Evoxac | 123603 | JPN | non-TD (10) | MD (10) | MDoff (7) |
| Adynovi | 86278357 | EU and JPN | non-TD (10) | CD (6) | MDon (10) |
| Gliolan | 123608 | EU and JPN | non-TD (10) | MD (10) | MDon (10) |
| Rinvoq | 58557659 | EU and JPN | non-TD (10) | CD (10) | MDoff (8) |
| Erizas | 15159004 | JPN | non-TD (10) | MD (10) | MDon (8) |
| Diquas | 148196 | JPN | non-TD (10) | CD (7) | MDon (9) |
| Olanedine | 67027685 | JPN | non-TD (10) | MD (10) | MDon (10) |
| Orapenem | 9892071 | JPN | non-TD (10) | MD (10) | MDon (8) |
| Xofluza | 124081896 | EU and JPN | non-TD (10) | CD (10) | MDon (7) |
| Intuniv | 71401 | EU and JPN | non-TD (10) | MD (10) | MDoff (8) |
| Tarceva | 176871 | EU and JPN | non-TD (10) | CD (10) | MDon (7) |
| Ninlaro | 56844015 | EU and JPN | non-TD (10) | MD (6) | MDon (10) |
| Mundesine | 135449327 | JPN | non-TD (10) | MD (8) | MDon (10) |
| Selincro | 5284594 | EU and JPN | non-TD (10) | MD (10) | MDoff (7) |
| Prevymis | 45138674 | EU and JPN | non-TD (10) | CD (8) | MDoff (6) |
| Lokelma | 91799284 | EU and JPN | non-TD (8) | MD (10) | MDon (10) |
| Revolade | 135449332 | EU and JPN | non-TD (10) | CD (10) | MDon (8) |
| Agrylin | 135413494 | JPN | non-TD (10) | MD (10) | MDoff (8) |
| Bretaris Genuair | 11519741 | EU and JPN | non-TD (10) | MD (10) | MDon (8) |
| Levitra | 135400189 | EU and JPN | non-TD (10) | CD (10) | MDon (9) |
| Isentress | 23668479 | EU and JPN | non-TD (10) | CD (10) | MDon (10) |
| Biktarvy | 129626368 | EU and JPN | non-TD (10) | MD (7) | MDon (9) |
| Cinacalcet Accordpharma | 156418 | EU and JPN | non-TD (10) | MD (10) | MDoff (8) |
| Actonel | 129628420 | JPN | non-TD (9) | MD (10) | MDon (10) |
| Recalbon | 23201029 | JPN | non-TD (10) | MD (10) | MDon (10) |
| Acofide | 6918406 | JPN | non-TD (10) | CD (10) | MDon (10) |
| Lonsurf | 9829639 | EU and JPN | non-TD (10) | CD (10) | MDon (10) |
| Corectim | 50914062 | JPN | non-TD (10) | MD (10) | MDon (6) |
| Alimta | 135413520 | JPN | non-TD (10) | CD (9) | MDon (9) |
| Sovriad | 118984468 | JPN | non-TD (10) | CD (7) | MDon (9) |
| Kiklin | 25212181 | JPN | non-TD (10) | MD (10) | MDon (8) |
| Treakisym | 77082 | JPN | non-TD (10) | MD (10) | MDoff (7) |
| Valixa | 135413534 | JPN | non-TD (10) | CD (7) | MDon (10) |
| Emend | 135413536 | EU and JPN | non-TD (10) | CD (10) | MDoff (6) |
| Proemend | 135413537 | JPN | non-TD (10) | MD (9) | MDon (9) |
| Vanflyta | 25184035 | JPN | non-TD (10) | CD (10) | MDon (9) |
| Equfina | 3038502 | JPN | non-TD (10) | CD (10) | MDon (10) |
| Oxarol | 6398761 | JPN | non-TD (10) | MD (10) | MDon (9) |
| Regtect | 155434 | JPN | non-TD (9) | MD (10) | MDon (10) |
| Unitalc | 26924 | JPN | TD (10) | MD (10) | MDon (10) |
| Revcovi | 121488177 | JPN | non-TD (10) | MD (10) | MDon (10) |
| Mephaquin | 65329 | JPN | non-TD (10) | CD (8) | MDoff (6) |
| Bronuck | 23663409 | JPN | non-TD (10) | MD (10) | MDon (6) |
| Voluven | 24846132 | JPN | TD (9) | MD (10) | MDon (9) |
| Beova | 44472635 | JPN | non-TD (10) | CD (10) | MDon (9) |

| | | | | | |
|---|---|---|---|---|---|
| Lasvic | 71528767 | JPN | non-TD (10) | CD (8) | MDon (8) |
| Olmetec | 130881 | JPN | non-TD (10) | CD (10) | MDon (9) |
| Tenelia | 53297474 | JPN | non-TD (10) | CD (10) | MDon (10) |
| Trevicta | 9852746 | EU and JPN | non-TD (10) | CD (10) | MDoff (7) |
| Narusus | 5462347 | JPN | non-TD (10) | MD (10) | MDoff (6) |
| Deferasirox Mylan | 214348 | EU and JPN | non-TD (10) | CD (10) | MDon (6) |
| Atripla | 464205 | EU and JPN | non-TD (10) | MD (10) | MDon (10) |
| Elplat | 9887053 | JPN | non-TD (10) | MD (10) | MDon (9) |
| Clozaril | 135398737 | JPN | non-TD (10) | MD (10) | MDoff (9) |
| Prodif | 214356 | JPN | non-TD (10) | CD (10) | MDon (10) |
| Valtrex | 135398741 | JPN | non-TD (10) | MD (7) | MDon (10) |
| Eybelis | 44230999 | JPN | non-TD (10) | CD (10) | MDon (9) |
| Mycobutin | 135398743 | JPN | non-TD (10) | MD (7) | MDon (9) |
| Jyseleca | 49831257 | EU and JPN | non-TD (10) | CD (10) | MDon (6) |
| Malarone | 9049 | JPN | non-TD (10) | MD (10) | MDon (5) |
| Olanzapine Apotex | 135398745 | EU and JPN | non-TD (10) | MD (10) | MDoff (10) |
| Zefnart | 3936 | JPN | non-TD (10) | MD (10) | MDoff (10) |
| Methapain | 14184 | JPN | non-TD (10) | MD (10) | MDoff (9) |
| Nicystagon | 23111531 | JPN | non-TD (10) | MD (10) | MDon (10) |
| Zinc-Tripentat | 164209 | JPN | non-TD (10) | MD (10) | MDon (10) |
| Elaspol | 23663985 | JPN | non-TD (10) | CD (9) | MDon (10) |
| Sinseron | 178038 | JPN | non-TD (10) | MD (8) | MDoff (7) |
| Minebro | 25052023 | JPN | non-TD (10) | CD (10) | MDon (7) |
| Enaroy | 50899324 | JPN | non-TD (10) | CD (10) | MDon (9) |
| Singulair | 23663996 | JPN | non-TD (10) | CD (7) | MDon (7) |
| Vitrakvi | 46188928 | EU and JPN | non-TD (10) | CD (10) | MDon (7) |
| Uriadec | 5288320 | JPN | non-TD (10) | MD (10) | MDon (8) |
| Hornel | 5282190 | JPN | non-TD (10) | MD (10) | MDon (8) |
| Takecab | 45375887 | JPN | non-TD (10) | CD (10) | MDon (10) |
| G-Lasta | 70683024 | JPN | non-TD (10) | MD (5) | MDon (10) |
| Parmodia | 11526038 | JPN | non-TD (10) | CD (10) | MDon (6) |
| Laserphyrin | 5488036 | JPN | non-TD (10) | MD (7) | MDon (10) |
| Benzodine | 65959 | JPN | non-TD (10) | MD (9) | MDoff (9) |
| Ditripentat-Cal | 25513 | JPN | non-TD (10) | MD (10) | MDon (10) |
| Lixiana | 25022378 | JPN | non-TD (10) | CD (10) | MDon (8) |
| Riona | 22178220 | JPN | non-TD (6) | MD (10) | MDon (10) |
| Seebri | 11693 | JPN | non-TD (10) | MD (10) | MDoff (7) |
| Nasonex | 441336 | JPN | non-TD (10) | MD (10) | MDon (9) |
| Acecide | 6585 | JPN | TD (6) | MD (10) | MDon (9) |
| Imusera, Gilenya | 107969 | JPN | non-TD (10) | MD (10) | MDon (7) |
| Precedex | 6918081 | JPN | non-TD (10) | MD (10) | MDoff (9) |
| Nexium Control | 130565 | EU | non-TD (8) | MD (6) | MDon (9) |
| Rasilez HCT | 51033608 | EU | non-TD (10) | CD (10) | MDon (8) |
| Dafiro HCT | 51033609 | EU | non-TD (10) | MD (6) | MDon (8) |
| Rizmoic | 56837137 | EU | non-TD (10) | CD (10) | MDon (10) |
| Tandemact | 11787804 | EU | non-TD (10) | MD (5) | MDon (9) |
| Vedrop | 53315106 | EU | non-TD (10) | MD (7) | MDon (5) |
| Tractocile | 5311010 | EU | non-TD (10) | MD (9) | MDon (10) |
| Arsenic trioxide medac | 14888 | EU | TD (10) | MD (10) | MDon (8) |
| Evotaz | 86583336 | EU | non-TD (10) | MD (6) | MDon (8) |
| DuoTrav | 16034857 | EU | non-TD (10) | MD (7) | MDon (9) |
| Ontozry | 11962412 | EU | non-TD (10) | MD (10) | MDon (8) |
| Xelevia | 11591741 | EU | non-TD (10) | CD (10) | MDon (10) |
| Feraccru | 169535 | EU | non-TD (8) | MD (10) | MDon (10) |
| Vokanamet | 53386314 | EU | non-TD (10) | CD (6) | MDon (10) |
| Quadramet | 56841803 | EU | non-TD (10) | MD (10) | MDon (10) |

| | | | | | |
|---|---|---|---|---|---|
| Fetcroja | 139033164 | EU | non-TD (10) | MD (9) | MDon (8) |
| Vabomere | 86298703 | EU | non-TD (10) | MD (10) | MDon (9) |
| Savene | 6918223 | EU | non-TD (10) | MD (10) | MDon (6) |
| IntronA | 71306834 | EU | non-TD (9) | CD (10) | MDon (7) |
| Trelegy Ellipta | 122403924 | EU | non-TD (10) | MD (10) | MDon (8) |
| Sogroya | 129894493 | EU | non-TD (10) | MD (7) | MDon (10) |
| Lamivudine | 160352 | EU | non-TD (10) | CD (10) | MDon (9) |
| Gencebok | 6241 | EU | non-TD (10) | CD (9) | MDon (10) |
| SomaKit TOC | 71661158 | EU | non-TD (10) | MD (10) | MDon (10) |
| Raxone | 3686 | EU | non-TD (10) | MD (10) | MDoff (6) |
| Zubsolv | 11274356 | EU | non-TD (10) | MD (10) | MDon (10) |
| Raloxifene Teva | 54900 | EU | non-TD (10) | MD (10) | MDon (8) |
| Stayveer | 185462 | EU | non-TD (10) | CD (10) | MDon (10) |
| Cholestagel | 60208760 | EU | non-TD (10) | MD (10) | MDon (7) |
| Qtern | 124219523 | EU | TD (9) | MD (8) | MDon (10) |
| Fareston | 3005572 | EU | non-TD (10) | MD (10) | MDon (10) |
| Mirvaso | 54405 | EU | non-TD (10) | MD (5) | MDon (9) |
| Vyxeos | 11422859 | EU | non-TD (10) | MD (6) | MDon (10) |
| Pioglitazone Teva | 60560 | EU | non-TD (10) | CD (10) | MDoff (7) |
| Omidria | 56950933 | EU | non-TD (10) | MD (8) | MDon (8) |
| Javlor | 6918295 | EU | non-TD (10) | MD (8) | MDon (9) |
| Taxespira | 148123 | EU | non-TD (10) | MD (8) | MDon (9) |
| Palonosetron Hospira | 6918303 | EU | non-TD (10) | MD (10) | MDoff (8) |
| Kivexa | 5273759 | EU | non-TD (10) | CD (10) | MDon (9) |
| Clopidogrel BGR | 115366 | EU | non-TD (10) | MD (10) | MDon (8) |
| Revinty Ellipta | 71306415 | EU | non-TD (10) | MD (10) | MDon (9) |
| Clopidogrel | 9938610 | EU | non-TD (10) | MD (10) | MDon (9) |
| Clopidogrel Apotex | 9847991 | EU | non-TD (10) | MD (10) | MDon (9) |
| EndolucinBeta | 71587001 | EU | non-TD (9) | MD (10) | MDon (8) |
| Resolor | 9870009 | EU | non-TD (10) | MD (8) | MDon (10) |
| Pemetrexed Krka | 135410875 | EU | non-TD (10) | CD (8) | MDon (9) |
| Pramipexole Accord | 166589 | EU | non-TD (10) | MD (10) | MDon (6) |
| Besremi | 86278347 | EU | non-TD (10) | MD (10) | MDon (10) |
| Dutrebis | 73386700 | EU | non-TD (10) | CD (10) | MDon (10) |
| Clopidogrel HCS | 9798860 | EU | non-TD (10) | MD (10) | MDoff (9) |
| Descovy | 90469070 | EU | non-TD (10) | CD (9) | MDon (10) |
| Firdapse | 9920716 | EU | non-TD (5) | MD (10) | MDon (10) |
| Jivi | 86278352 | EU | non-TD (10) | MD (10) | MDon (10) |
| Cresemba | 72196309 | EU | non-TD (10) | MD (7) | MDon (8) |
| Tolucombi, Actelsar HCT | 216293 | EU | non-TD (10) | CD (10) | MDon (9) |
| Mysimba | 11556075 | EU | non-TD (10) | MD (10) | MDon (10) |
| Pemetrexed Accord | 135564527 | EU | non-TD (10) | CD (8) | MDon (8) |
| Zoledronic Acid Accord | 121586 | EU | non-TD (10) | MD (10) | MDon (10) |
| Tookad | 129629432 | EU | non-TD (10) | MD (7) | MDon (10) |
| Anagrelide Mylan | 135409400 | EU | non-TD (10) | MD (10) | MDoff (8) |
| Caspofungin Accord | 6850808 | EU | non-TD (10) | MD (10) | MDon (10) |
| Kuvan | 135398654 | EU | non-TD (10) | MD (10) | MDon (10) |
| Fosavance | 23682309 | EU | non-TD (10) | MD (10) | MDon (10) |
| Cufence | 71433 | EU | non-TD (7) | MD (10) | MDon (9) |
| Cuprior | 71434 | EU | non-TD (6) | MD (10) | MDon (9) |
| Cayston | 5459211 | EU | non-TD (10) | MD (10) | MDon (9) |
| Evrysdi | 118513932 | EU | non-TD (10) | CD (10) | MDon (6) |
| Orkambi | 71494926 | EU | non-TD (10) | CD (10) | MDon (8) |
| Rasagiline Mylan | 71550735 | EU | non-TD (10) | MD (10) | MDon (10) |
| Veltassa | 86580497 | EU | non-TD (10) | MD (10) | MDon (9) |
| Zoely | 9895694 | EU | non-TD (10) | MD (10) | MDon (10) |

| | | | | | |
|---|---|---|---|---|---|
| Sildenafil Actavis | 135413523 | EU | non-TD (10) | CD (10) | MDon (10) |
| Oslif Breezhaler, | 9827599 | EU | non-TD (10) | CD (10) | MDon (10) |
| Roclanda | 118984471 | EU | non-TD (10) | MD (7) | MDon (10) |
| Talzenna | 135565082 | EU | non-TD (10) | CD (10) | MDon (7) |
| Triumeq | 54736666 | EU | non-TD (10) | CD (10) | MDon (9) |
| Duzallo | 137083676 | EU | non-TD (10) | CD (10) | MDon (10) |
| Potactasol | 60699 | EU | non-TD (10) | CD (10) | MDon (10) |
| Ivemend | 135413538 | EU | non-TD (10) | CD (10) | MDon (10) |
| Eucreas | 121488163 | EU | non-TD (10) | MD (8) | MDon (10) |
| Eptifibatide Accord | 448812 | EU | non-TD (10) | MD (10) | MDon (10) |
| Lamzede | 32051 | EU | TD (10) | MD (10) | MDon (8) |
| Trepulmix | 23663413 | EU | non-TD (10) | MD (10) | MDon (10) |
| Brinavess | 9930048 | EU | non-TD (10) | MD (10) | MDon (5) |
| Symkevi | 72722243 | EU | non-TD (10) | CD (10) | MDon (10) |
| Aerinaze | 9891141 | EU | non-TD (10) | CD (8) | MDoff (7) |
| Numient | 104778 | EU | non-TD (10) | MD (10) | MDon (10) |
| Foscan | 60751 | EU | non-TD (10) | CD (10) | MDon (9) |
| Ganfort | 126480208 | EU | non-TD (7) | CD (10) | MDon (10) |
| Macimorelin Aeterna Zentaris | 71526737 | EU | non-TD (10) | CD (9) | MDon (10) |
| Karvezide | 9831761 | EU | non-TD (10) | CD (10) | MDon (9) |
| Lopinavir | 11979606 | EU | non-TD (10) | MD (10) | MDon (9) |
| Ulipristal Acetate Gedeon Richter | 130904 | EU | non-TD (10) | MD (10) | MDon (5) |
| Granpidam | 135398744 | EU | non-TD (10) | CD (10) | MDon (10) |
| Oncaspar | 436058 | EU | non-TD (7) | MD (10) | MDoff (9) |
| Mepact | 23663962 | EU | non-TD (10) | MD (10) | MDon (10) |
| Myocet | 50925403 | EU | non-TD (10) | MD (10) | MDon (10) |
| Wilzin | 2724192 | EU | TD (10) | MD (10) | MDon (10) |
| Iblias | 104815 | EU | TD (10) | MD (10) | MDon (8) |
| Cuprymina | 76972915 | EU | non-TD (10) | MD (10) | MDon (8) |
| Fexeric | 61300 | EU | TD (8) | MD (10) | MDon (10) |
| Ayvakyt | 118023034 | EU | non-TD (10) | CD (10) | MDon (10) |
| Juluca | 131801472 | EU | non-TD (10) | CD (10) | MDon (10) |
| Procysbi | 11958147 | EU | non-TD (10) | MD (10) | MDon (10) |
| Xermelo | 25181577 | EU | non-TD (10) | CD (10) | MDon (9) |
| Spectrila | 5460875 | EU | non-TD (10) | MD (10) | MDon (10) |
| Jentadueto | 46861711 | EU | non-TD (10) | CD (10) | MDon (10) |
| Sensitivity | | | 0.96 | 0.62 | 0.84 |

The list of Europe and Japan approved drugs was collected from KEGG DRUG database (https://www.genome.jp/kegg/drug/ ). After removing those appeared in our compound datasets (Figure 5) and duplicates, as well as those exceeding molecular weight of 1500 Daltons, a total of 256 drugs were collected, of which 49 were approved by both the European Union and Japan, as indicated. Sensitivity was calculated for the positive group, i.e. non-TD for the non-TD/TD predictions, MD for the MD/CD predictions, and MDon for the MDon/MDoff predictions. Number in parenthesis indicates the number of models predicting the class (five indicates absence of a majority prediction). TD, toxic compound; CD, drug in clinical trial; MD, drug approved by FDA; MDoff, drug withdrawn from the market; MDon, drug currently on the market; EU, European Union; JPN, Japan.

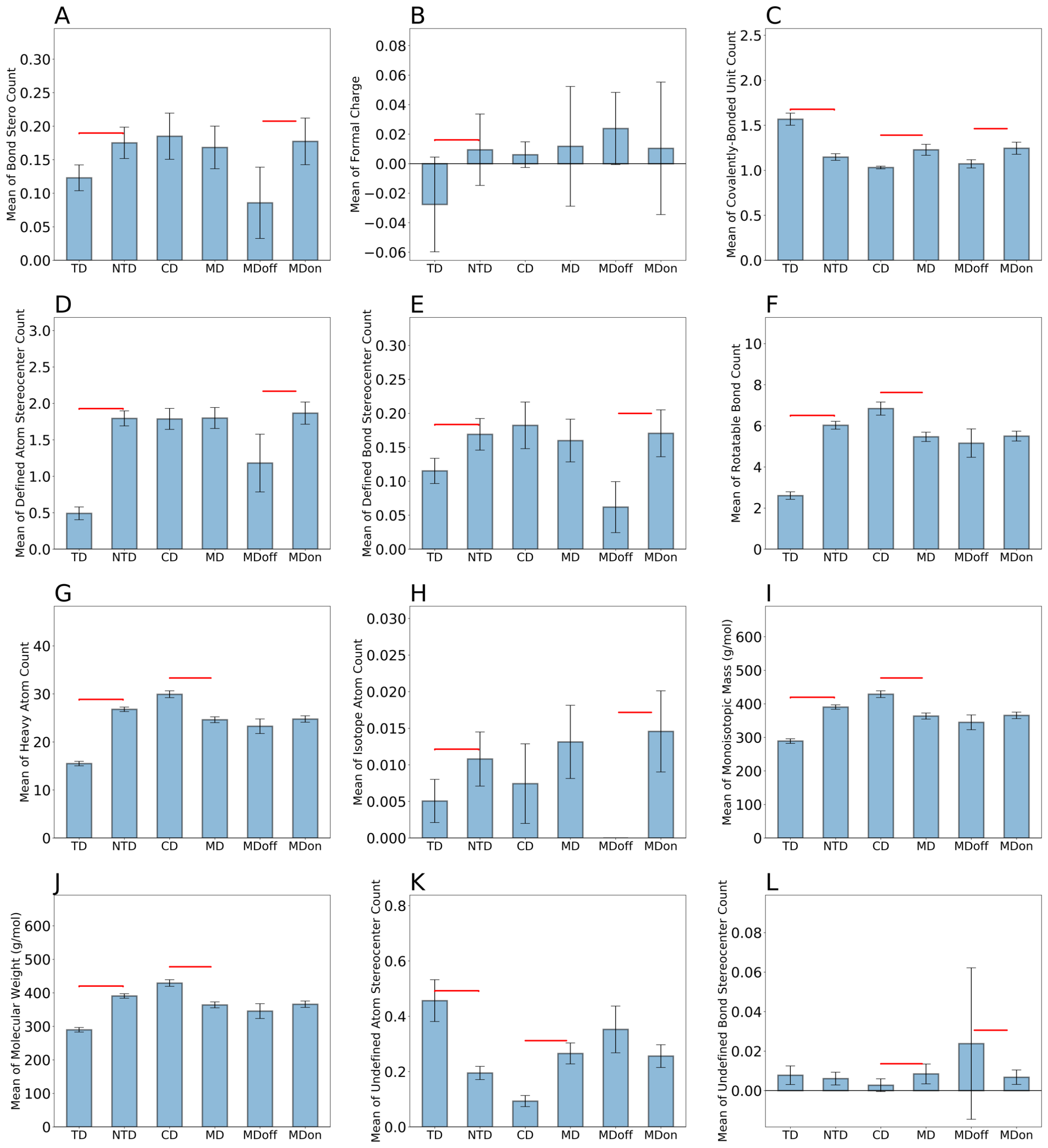


**Figure S1.** Physicochemical property comparison results between (1) TD vs. non-TD, (2) CD vs. MD, and (3) MDoff vs. MDon. Bar plots show calculated means of each property in each compound group. Difference was tested using one-tailed Student's t-test for each comparison. Error bar: 95% confidence interval, red significance bridges: p value of t test<0.05. NTD, nontoxic; TD, toxic; CD, in clinical trial; MD, approved by FDA; MDoff, withdrawn from the market; MDon, currently on the market.

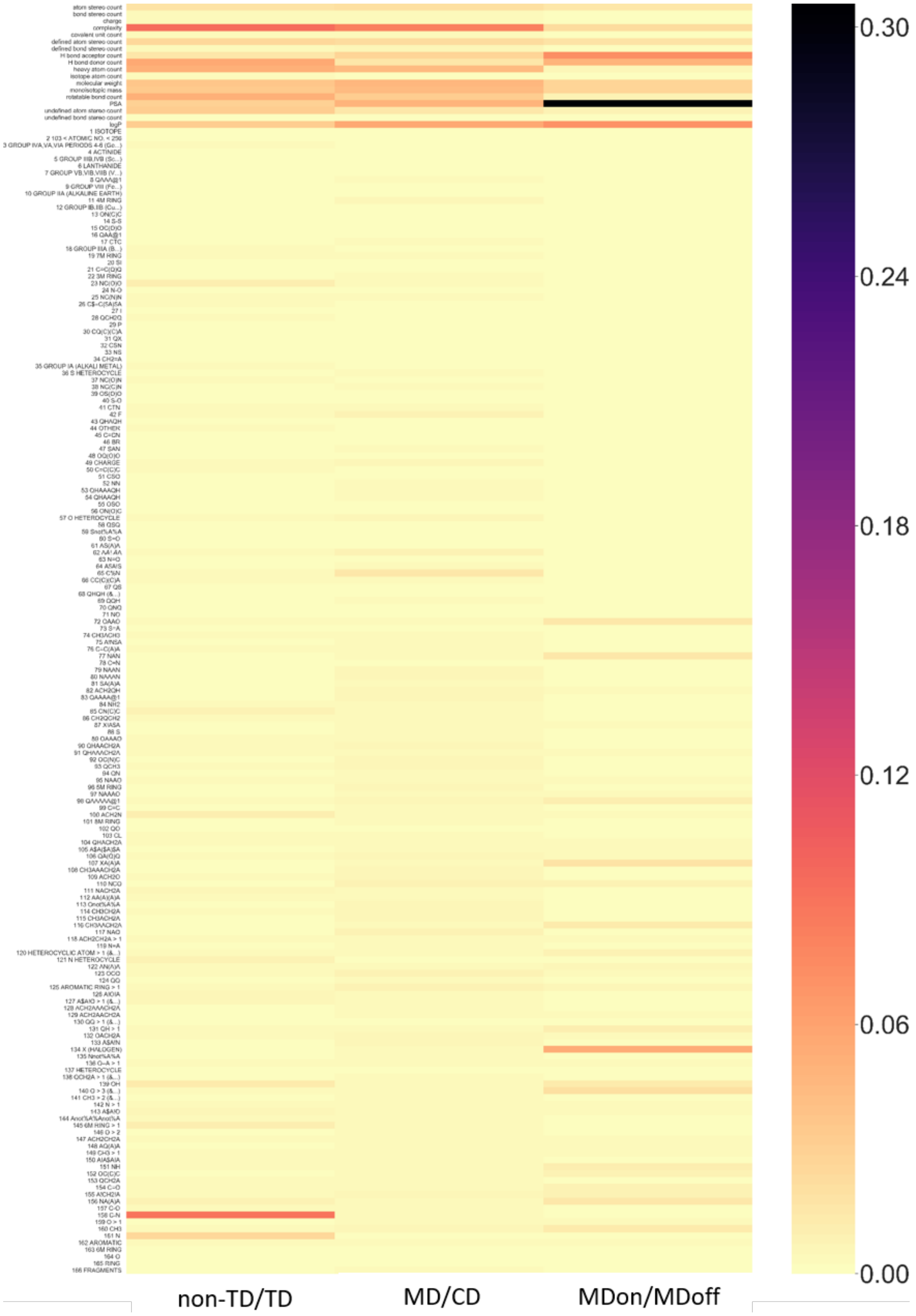


**Figure S2.** Heat map of feature importance for all features used by the machine learning models. To the right is the importance score bar (darker color indicates more importance). See Durant et al.'s report[6] and RDkit's[7] GitHub repository for the ID number and definition of the Molecular

Access System fingerprints. TD, toxic compound; CD, drug in clinical trial; MD, drug approved by FDA; MDoff, drug withdrawn from the market; MDon, drug currently on the market.

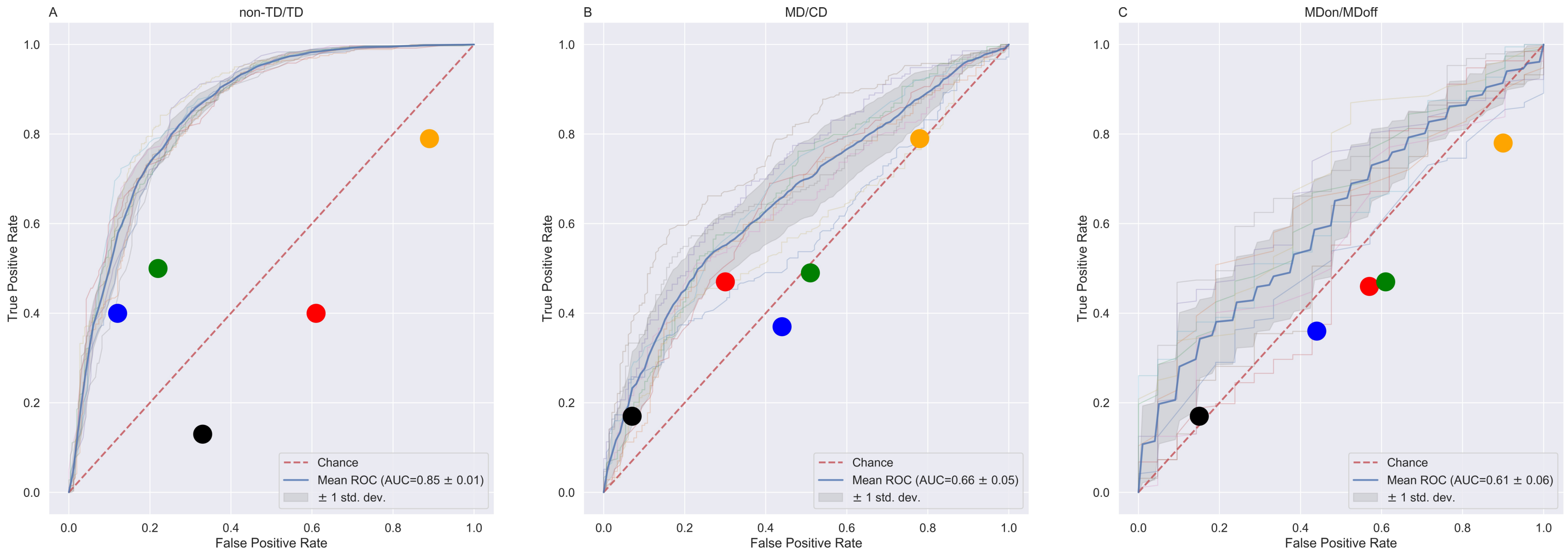


**Figure S3.** Model performance using only the 18 physicochemical properties. These were model's prediction results on test set compounds (see Materials and Methods) for the three classification comparisons of non-TD/TD, MD/CD, and MDon/MDoff. Upper panel: the ROC (receiver operating characteristic) curves for 10 independent models, each being a gray line and the blue line is their mean ROC, on the classification of (A) non-TD/TD, (B) MD/CD, and (C) MDon/MDoff. Color dots indicate the performance of Rule of Five (red), Ghose filter (blue), Veber filter (orange), Rule of Three (black), and Rapid Elimination of Swill (REOS) filter (green).

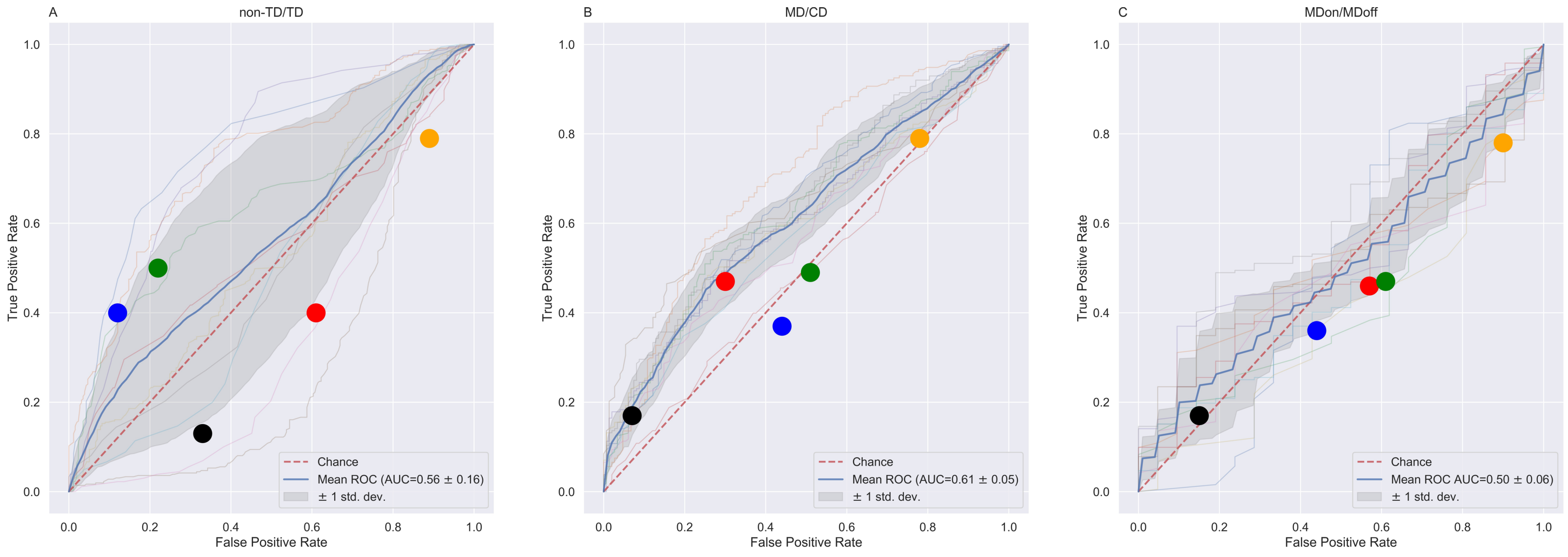


**Figure S4.** Model performance using only the Molecular Access System fingerprints. The model predictions were for test set compounds (see Materials and Methods) for the three classification comparisons (non-TD/TD, MD/CD, and MDon/MDoff). Upper panel: receiver operating characteristic (ROC) curves for 10 independent models (gray lines) and mean ROC (the blue lines) on the classification of (A) non-TD/TD, (B) MD/CD, and (C) MDon/MDoff. Color dots indicate performance on Rule of Five (red), Ghose filter (blue), Veber filter (orange), Rule of Three (black), and Rapid Elimination of Swill (REOS) filter (green).
